# Automatic detection of seizures within zebrafish larvae epilepsy models using single-channel electroencephalography

**DOI:** 10.1101/2024.08.18.608512

**Authors:** Connor Delaney, Qi Wei, Nathalia Peixoto

## Abstract

Researchers continue to pursue new drugs capable of treating intractable, or drug- resistant, epilepsy as a large number of patients do not see a reduction in the number of seizures from current treatments. To quicken the pace of drug research, zebrafish (*Danio rerio*) have been utilized as a model organism for testing anticonvulsant drugs before clinical trials. However, the lengthy task of labeling electroencephalography (EEG) data slows the pace of this line of research and limits its full potential. This study investigated the creation of automatic seizure detection algorithms for electroencephalogram data recorded from seizure-induced zebrafish to detect seizure, artifact, and neurotypical events. Four unique seizure detection algorithms were proposed and implemented based on k-nearest neighbor (KNN), support vector machine (SVM), and artificial neural network (ANN) classifiers. These four techniques were tested using the same input features and their results compared. The best-performing algorithm was identified as the Single Stage KNN with an 83.8% accuracy, followed by the Single Stage ANN with an 80.3% accuracy. The results indicate that a single-stage, multiclass classification architecture may be beneficial to automatically labeling epilepsy data, thus enhancing efficiency. Furthermore, the results for the algorithms which separate the multiclass classification into a series of binary classifications suggest that there are advantages in research and clinical settings to implement a detection algorithm that can delineate neurotypical and non-neurotypical data to assist with manual labeling.

## Introduction

Epilepsy is a term used to describe a broad collection of neurological disorders characterized by recurrent seizures. Despite anticonvulsant drugs being able to suppress seizures in many patients, approximately a third of individuals experience intractable epilepsy that remain refractory to interventions [1,2]. In patients with intractable epilepsy, electroencephalograms (EEG) are often recorded and used for planning surgical resections of the brain regions where the patient’s seizures originate. A large body of research has been committed to automatically detecting seizures from EEG recordings, as trained clinicians have the lengthy task of labeling patients’ EEG data for seizure occurrences. Automatic detection algorithms able to accurately detect periods when seizures occurred in patient recordings would allow clinicians to spend less time on this task and more time addressing other patient needs. Additionally, utilizing an automated detection system would remove operator-dependent errors and allow for repeatability across datasets.

Many different approaches have been taken to designing seizure detection algorithms for EEG data from humans, including ones based on signal filtering procedures, linear discriminant classifiers, k-nearest neighbor classifiers (KNN), decision trees, spike detectors, support vector machines (SVM), and artificial neural networks (ANNs) [3,4]. Typically, these studies employ a relatively large number of electrodes with features from all these channels simultaneously to detect seizures.

Concurrently, researchers continue to explore genetic links to epilepsy and its severity, as well as developing new drugs that can treat epilepsy resistant to on-market medications. As human trials incur more hurdles for drug trials, the epilepsy research community has adopted several animal models to aid in the speed of drug development [5]. One such animal model is the zebrafish (Danio rerio), which has several ideal qualities as an animal model, such as 70% homology with human genes, a fully sequenced genome, and a high fecundity rate [6,7,8].

Furthermore, both adult and larval zebrafish have been used to study epilepsy. Larval zebrafish are sometimes favored due to their translucent bodies, while adults are larger thus allowing for more electrodes to be utilized when recording electrophysiological signals. Because of these benefits of using zebrafish as a model organism, there is a small body of literature on seizure detection algorithms, specifically using data collected from zebrafish. Predominantly, these seizure detection methods include video-based methods [9]. These methods often utilize videos that image a certain biomarker of nervous system behavior, such as calcium. While calcium imaging methods are solid tools for monitoring the seizure activity of translucent zebrafish larvae, they require sophisticated and expensive experimental setups. Additionally, relating calcium imaging to the EEG recordings from zebrafish larvae and more broadly EEG from humans presents a challenging task for researchers as these imaging tools have very different spatial and temporal recording properties [10]. EEG presents an ideal modality for seizure detection within zebrafish models as it is less costly than imaging, has a dramatically higher time resolution, and is used to monitor human epilepsy patients. This has led to a specialized field of research forming around studying zebrafish epilepsy models with EEG. Custom recording hardware and specialized techniques have been developed by researchers to capture EEG from zebrafish including multichannel arrays for adult zebrafish, techniques to immobilize zebrafish throughout long recording sessions, cheap 3D printed alternatives to traditional recording electrodes, and systems able to record from multiple zebrafish simultaneously [11–16].

Additionally, EEG recordings from seizure-induced zebrafish models have been utilized to verify the efficacy of anticonvulsant drugs with similar capabilities to calcium imaging [17]. However, there is still much work to be done regarding improving the accuracy of seizure detection using EEG with zebrafish models. Hong *et al.* (2016) developed a simple algorithm taking the time derivative and cross-correlation of recorded signals to count seizures over lengthy recordings with multichannel recordings [14]. Hunyadi *et al.* (2017) were able to take pre-segmented and labeled data containing seizures, artifacts, and “dubious” data and classify it leveraging k-nearest neighbors and support vector machine algorithms [18]. The results from this study demonstrated the possibility of high specificity classification of non-neurotypical events. Still, they relied upon manual labeling and segmentation of these out of the original body of their recordings. This is less than ideal as it does not address the primary reason for implementing an automatic seizure detection algorithm: the large time requirement for human technicians to label the data.

Although detecting and predicting seizures has been researched for some time now, the need to improve upon existing algorithms for this task still exists. This is especially the case when the amount of available data is limited, such as in low channel count EEG recordings like the ones used often with zebrafish models. Therefore, it was proposed that single-channel EEG recordings in larval zebrafish be investigated as a model of epilepsy in humans and that different seizure detection methods be compared.

We employed four distinct detection algorithms were employed and compared against one another. A KNN classifier was used as a simple, data-driven algorithm that could handle multiclass detection. A multiclass, fully connected feed-forward ANN was employed as the complex detection algorithm to compare against the KNN classifier. We referred to these two algorithms as “Single Stage” since they approached the detection problem using a single classifier capable of multiclass classification. “Two Stage” detection algorithms were also employed to split the problem into two binary classification problems (neurotypical vs. non-neurotypical and seizure vs. artifact). The first of these two-stage detection algorithms was a pair of Support Vector Machine (SVM) classifiers. The second two-stage detection algorithm was a pair of ANN binary classifiers. A diagram presenting the structure of the single- and two-stage algorithms is presented in Fig 1. All these detection algorithms were designed with the ability to discriminate between neurotypical, ictal (seizure), and artifact-corrupted data. This, in theory, removes the need for lengthy data labeling sessions by technicians if implemented in practice.

**Fig 1.**
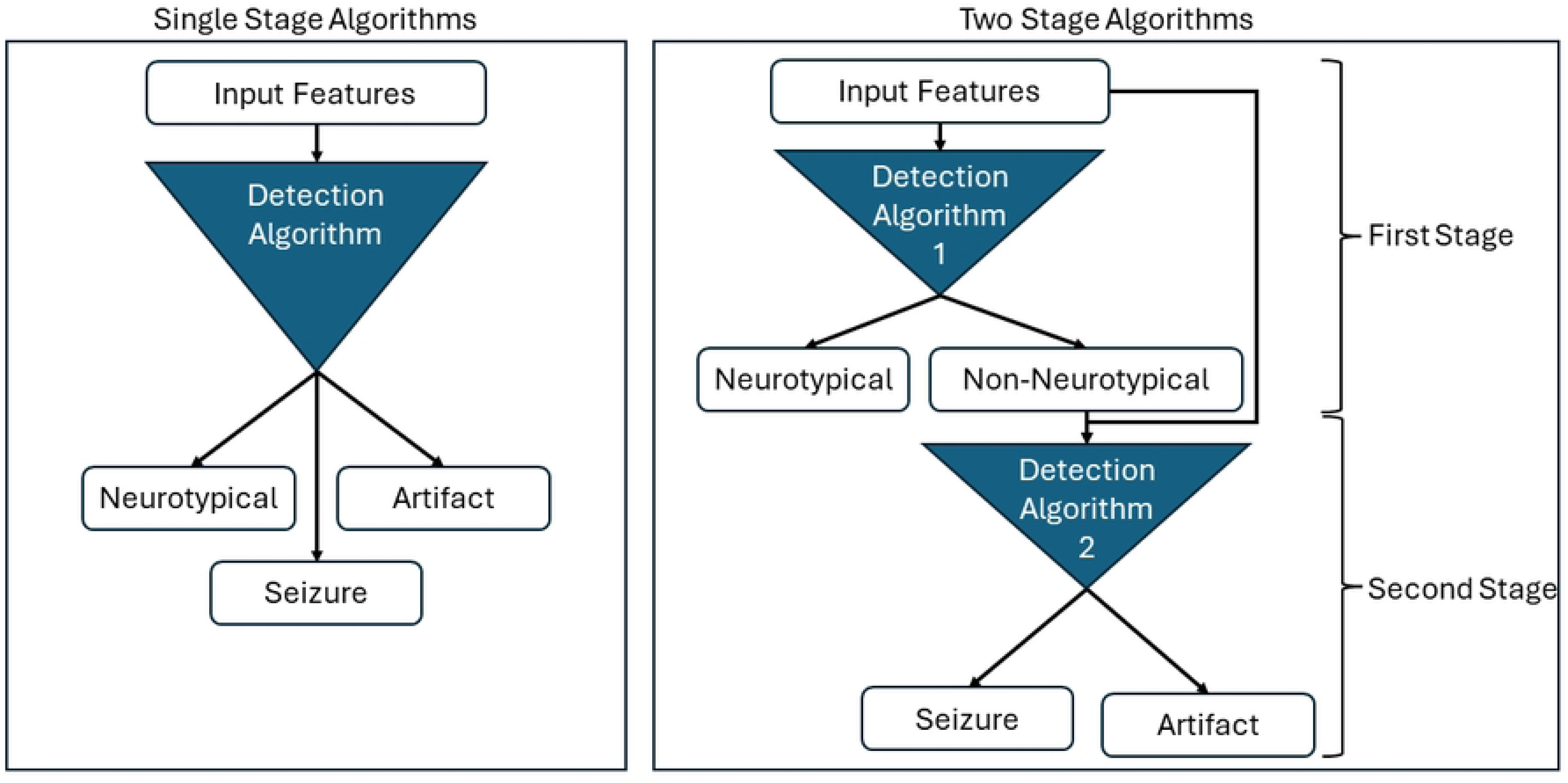
Diagram Comparing Single Stage and Two Stage Detection Algorithms. The single stage detection algorithms were multi-class classifiers which take the input features and assign one class (neurotypical, seizure, or artifact) to the data. The two stage detection algorithms were comprised of two separate binary classifiers. The first of these classifiers labels the data as neurotypical or non-neurotypical. If data was labeled as non-neurotypical, then the input features were passed to the second classifier. The second classifier assigned the non-neurotypical data a label of seizure or artifact.

## Methods

### Animals and Data Collection

We used previously recorded data collected from specimens as described in Xing *et al*.

2018 [19]. Details of animal care for this study are thoroughly described in the same publication. Animal procedures and care were approved by the Ethics Committee on the Use of Animals (CEUA \#4660-1 and \#4785-1) of UNICAMP, Campinas, Brazil.

A pentylenetetrazol (PTZ) induced seizure method was used to model epilepsy within zebrafish larvae [20,21,22]. EEG was recorded from the larvae between 5 to 16 days post fertilization with stainless steel electrodes with 125-micron diameter. Larvae were anesthetized in .002% tricaine and 10 micromolar d-tubocurarine and then immobilized at a 1.2% concentration of low-melting point agarose. The electrodes were placed within the tectum opticum with a binocular loupe and a triaxial micro-manipulator. The electrical signal was pre-amplified using a C3313/RHD2216 and amplified by a C3100/RHD2000 (both devices from Intan Technologies LLC) for a final gain of 1000. The single-channel recordings were collected at a 10 kHz sampling rate. Before each recording session, 30 mM PTZ solution was added to the bath, and then recording sessions were performed with the given larva. The data presented in this present work consists of 20 of these recording sessions for a total duration of 243 minutes of raw data, which have had the time stamps of seizures and artifacts within them labeled by trained technicians using the labeling guidelines as presented in Hunyadi *et al.* (2017) [18].

### Data Preprocessing

All data preprocessing and feature extraction were performed in MATLAB. The signals were preprocessed by downsampling from 10 kHz to 1 kHz. They were then bandpass filtered from 1 to 50Hz.

The detection algorithms were trained and created on two distinct versions of the data, each using a unique windowing procedure. The first procedure used windows of the data that were 4 seconds in length with a 75% overlap. The second type used 1-second duration windows with 50% overlap. If a window contained any data labeled as a seizure or artifact, it was assigned the same label for classification. If a window contained no time labeled as a seizure or artifact, it was assigned the neurotypical behavior label. Each window was then normalized by subtracting the mean amplitude of the window signal and dividing it by the standard deviation of the entire recording. The maximum absolute amplitude, root mean square, sample entropy, and signal energy were extracted from each time window. Next, a Kaiser window was applied to each window to reduce the Gibbs phenomenon before a Fourier transform was taken to convert the signal to the frequency domain. The features extracted from the frequency domain signal included the frequency with the maximum magnitude, average power across all frequencies, delta band power, theta band power, alpha band power, beta band power, and gamma band power. All features are summarized in Table 1. An example window of data is displayed in Fig 2. The individual features for all windows were combined into one data set which was then split for training and testing each algorithm.

**Fig 2.**
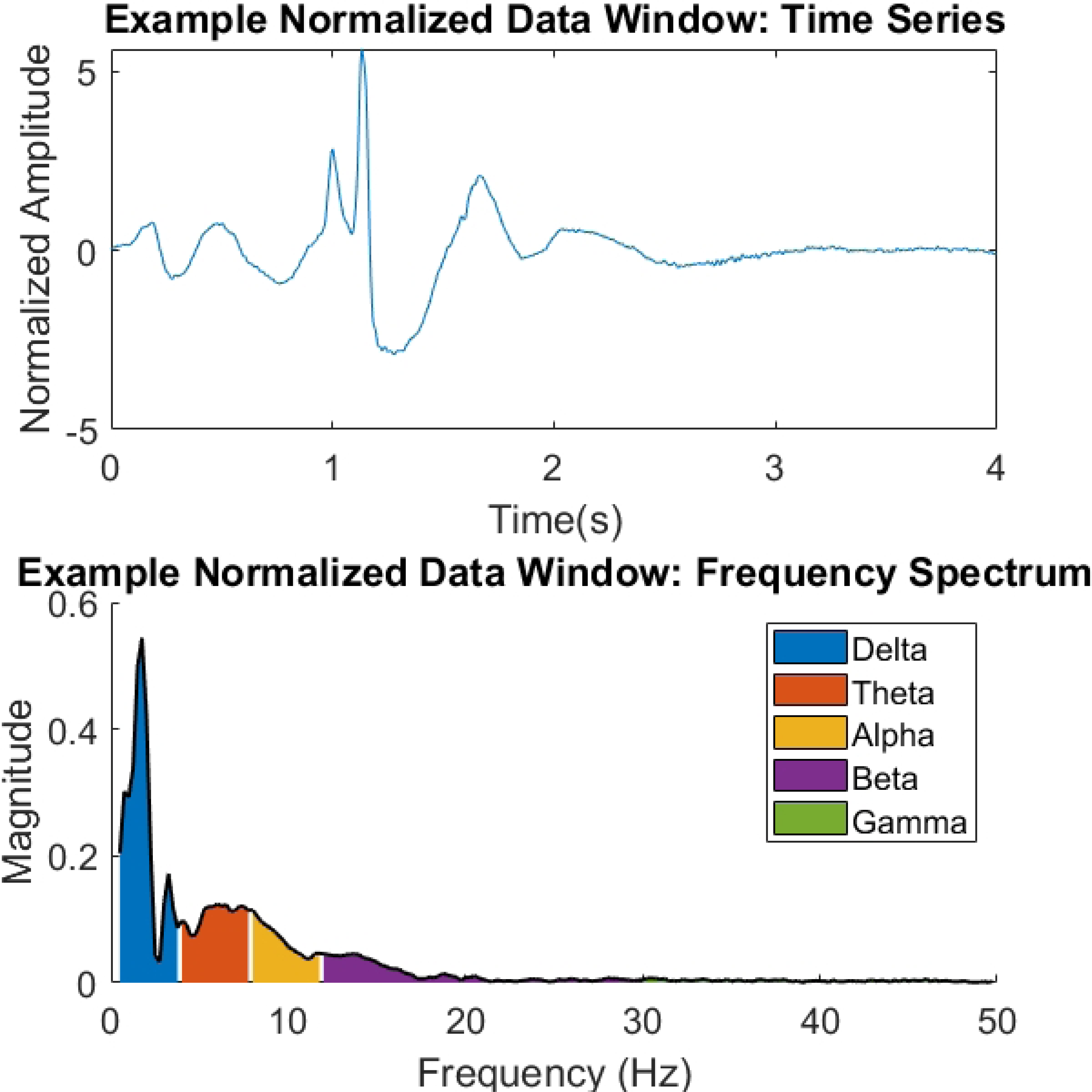
Example of Data Window. An example of a window of data containing a seizure and its frequency domain representation.

**Table 1.** List of Detection Algorithm Features.

| Feature Name | Description |
| --- | --- |
| Time Absolute Maximum | The maximum absolute value of amplitude from the time series. |
| Time Variance | The variance of the amplitude from the time series. |
| Time Root Mean Square | The root mean square value of amplitude from the time series. |
| Time Sample Entropy | The sample entropy of the time series. Sample Entropy was calculated as follows:<br>$\sum_{i=1}^N (X_i * \log X_i^2)$ |
| Time Energy | The signal energy of the time series. Signal energy was calculated as:<br>$\sum_{i=1}^N (x_i^2)$ |
| Peak Frequency | The frequency with the maximum magnitude from the frequency domain view of the window. |
| Average Power | The average magnitude across all frequency bins of the frequency domain view of the window. |
| $\delta$ Band Power | The power of the frequency band is from .5 to 4Hz of the frequency domain view of the window. |
| $\theta$ Band Power | The power of the frequency band is from 4 to 8Hz for the frequency domain view of the window. |
| $\alpha$ Band Power | The power of the frequency band is from 8 to 12Hz in the frequency domain view of the window. |
| $\beta$ Band Power | The power of the frequency band is from 12 to 30Hz in the frequency domain view of the window. |
| $\gamma$ Band Power | The power of the frequency band is from 30 to 50Hz in the frequency domain view of the window. |

All features were extracted from downsampled, filtered, and normalized data.

### Seizure Detection Algorithms

The four seizure detection algorithms based upon standard classification techniques were all implemented with the same 10-fold cross-validation procedure but with different organizations of the data between the single- and two-stage algorithms. Each fold consisted of 10% of the windows in the dataset being reserved for validation data and the remaining data being used as the training data. The training and validation subsets of data were both class-balanced. The data assigned to the validation set in one fold was not assigned to the validation set in another. Table 2 presents the size of the 4-second window dataset, and Table 3 presents the size of the 1-second window dataset.

**Table 2.** Description of the 4 Second Window Data.

| Data Set Name | Description | Size of Training Data per Fold | Size of Validation Data per Fold | Algorithms using Dataset |
| --- | --- | --- | --- | --- |
| Single Stage Dataset | Class-balanced dataset with labels for seizures, artifacts, and neurotypical data. | 3483 | 387 | Single Stage KNN, Single Stage ANN |
| Two Stage Dataset – First Stage | Class-balanced dataset with labels for neurotypical and non-neurotypical data (seizure and artifact). | 4662 | 518 | Two Stage SVM (First Stage), Two Stage ANN (First Stage) |
| Two Stage Dataset – Second Stage | Class-balanced dataset with labels for seizure and artifact data. This dataset only included data from seizures and artifacts. | 2322 | 258 | Two Stage SVM (Second Stage), Two Stage ANN (Second Stage) |

Each training and validation data set was class balanced within each cross-validation fold.

**Table 3.** Description of 1 Second Window Data.

| Data Set Name | Description | Size of Training Data per Fold | Size of Validation Data per Fold | Algorithms using Dataset |
| --- | --- | --- | --- | --- |
| Single Stage Dataset | Class-balanced dataset with labels for seizures, artifacts, and neurotypical data. | 4590 | 510 | Single Stage KNN, Single Stage ANN |
| Two Stage Dataset – First Stage | Class-balanced dataset with labels for neurotypical and non-neurotypical data (seizure and artifact) | 6138 | 682 | Two Stage SVM (First Stage), Two Stage ANN (First Stage) |
| Two Stage Dataset – Second Stage | Class-balanced dataset with labels for seizure and artifact data. This dataset only included data from seizures and artifacts. | 3060 | 340 | Two Stage SVM (Second Stage), Two Stage ANN (Second Stage) |

Each set of training and validation data was class balanced within each cross-validation fold.

The first detection method consisted of a k-nearest neighbors classifier, referred to as the “Single Stage KNN” algorithm. The training data was used as the database constituting the neighbors. The standardized Euclidean distance was used as the distance metric to measure the similarity between two data points. This means that each of the windows’ feature variables was scaled by the standard deviation of the respective feature in the training data. K values ranging from 1 to 100 were tested on each fold, and the K value producing the maximal average accuracy across the ten cross-validation folds was selected as the optimal K value used on the test data to evaluate the algorithm. In the case of ties for the maximal average accuracy of given k-values, the smallest k-value greater than one was selected. This method was implemented and evaluated in MATLAB.

The second detection method comprised a fully connected feed-forward artificial neural network. This detection algorithm was referred to as the “Single Stage ANN.” The network’s inputs were the standardized features of each window, and the output was three probabilities, each representing the odds of the input window belonging to one of the three classes (seizure, neurotypical, and artifact). Fully connected linear layers using horizontal tangent activation functions followed by a dropout made up of each layer. The network consisted of 3 layers and is presented in Fig 3. The network’s hyperparameters for training are contained in Table 4, and a batch training procedure was utilized. After training, class probabilities were converted into class labels by selecting the class with the highest probability. This seizure detection method was implemented and evaluated in Python using PyTorch.

**Fig 3.**
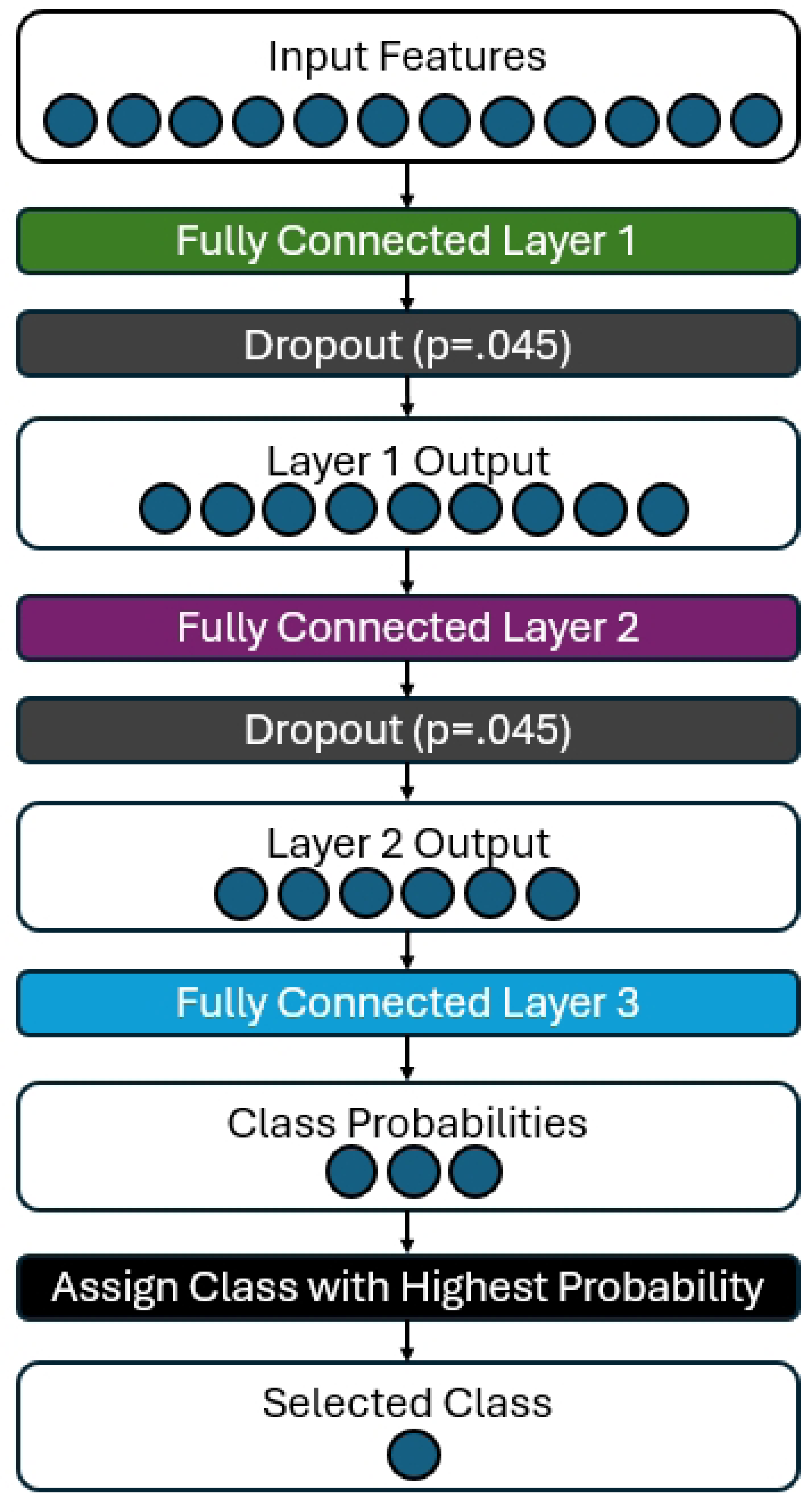
Diagram of the Single Stage ANN architecture used for seizure detection. The 12 features from the original dataset acted as the input to the first layer. Each layer consisted of a fully connected linear layer and dropout. Horizontal tangent activation functions were used with the fully connected linear layers. Dropout was performed with a 4.5% chance of any weight in the layer being set to zero for that epoch. The network output was the probability of the input being from the three classes (seizure, neurotypical, and artifact). The class with the highest probability was then assigned to the window.

**Table 4.** Seizure Detection Artificial Neural Network Training Hyperparameters.

| Feature Name | Description |
| --- | --- |
| Number of Epochs | 30000 for Single Stage ANN.<br>30000 for Two Stage ANN – Second Stage.<br>7500 for Two Stage ANN - First Stage. |
| Learning Rate | .00225 |
| Loss Function | Mean Square Error |
| Objective Optimization Algorithm | Adam |
| Adam $\beta_1$ | .5 |
| Adam $\beta_2$ | .999 |
| Weight Initialization | Golrot Normal Distribution Initialization |
| Bias Initialization | All bias values were initialized to .01 |

These hyperparameters were used for both the four and one-second window duration datasets.

The third seizure detection algorithm was the “Two Stage SVM.” This detection method contained a pair of C-SVM binary classifiers that worked sequentially on an input after training. The first stage classifier classified the input as neurotypical or non-neurotypical (seizures and artifacts). The second stage classifier then classified the input as a seizure or artifact. In practice, when the first stage classifies an input window as non-neurotypical, the window is then passed along to the second stage, which attempts to delineate artifacts from seizures. Only windows detected as non-neurotypical were passed from the first stage to the second stage.

During the design of the two-stage SVM, several kernels were compared: linear, polynomial, radial basis function (RBF), and sigmoid. Furthermore, gamma values of 0.1 to 1 and C values of 0.1 to 2 were tested as well. We report only the results for the best-performing Two-Stage SVM kernel implementation.

The fourth seizure detection algorithm consisted of a pair of binary classification fully connected feed-forward ANNs. This detection algorithm was referred to as the “Two Stage ANN.” Similar to the Single Stage ANN, this network used the standardized features of each window as the input. The output for the first stage was the probability that the input window was neurotypical or non-neurotypical (seizure or artifact). The output from the second stage was the probability that the input window was from seizure or artifact corrupted data. Both networks in the pair were trained separately and then used in conjunction for combined validation. Validation data was first input to the first stage network, and if it were classified as non-neurotypical, it would be sent to the second stage classifier for final classification. Fully connected linear layers using horizontal tangent activation functions followed by dropout were used for the first layer, and the second layer produced the output. A diagram of a single stage of the network is presented in Fig 4. The network’s hyperparameters for training are presented in Table 4, and a batch training procedure was utilized as with the Single Stage ANN. After training, class probabilities were converted into class labels by selecting the class with the highest probability. This seizure detection method was implemented and evaluated in Python using PyTorch. A diagram of the complete study methodology is contained in Fig 5.

**Fig 4.**
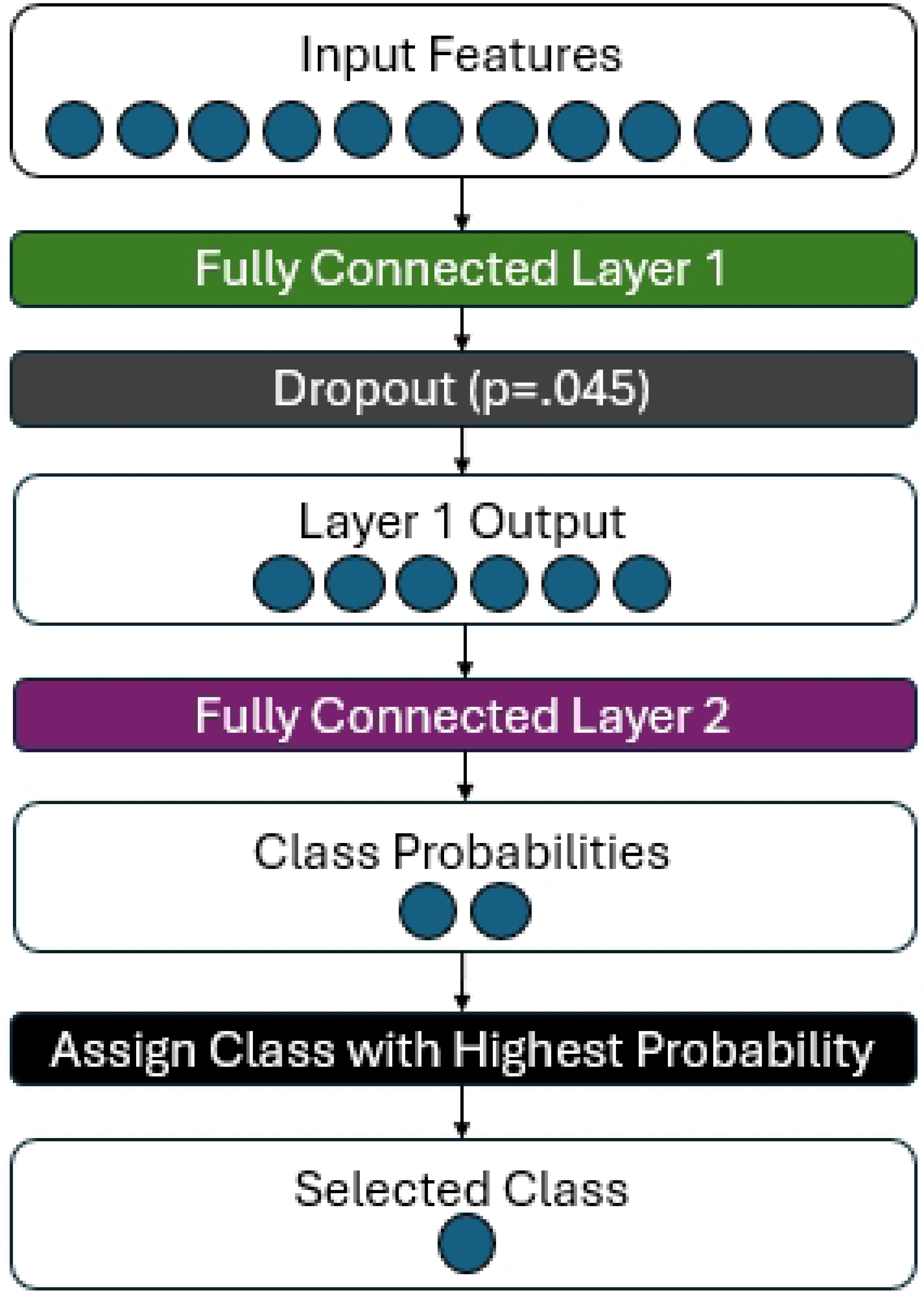
Diagram of the Two Stage ANN architecture used for seizure detection. The 12 features from the original dataset acted as the input to the first layer. Each layer consisted of a fully connected linear layer and dropout. Horizontal tangent activation functions were used with the fully connected linear layers. Dropout was performed with a 4.5% chance of any weight in the layer being set to zero for that epoch. The network output was the probability of the input being from the two classes being considered for that stage (neurotypical and non-neurotypical for the first stage; seizure and artifact for the second stage). The class with the highest probability was then assigned to the window. After training, if the output of the first stage was non-neurotypical then the window was passed to the second stage network for final classification.

**Fig 5.**
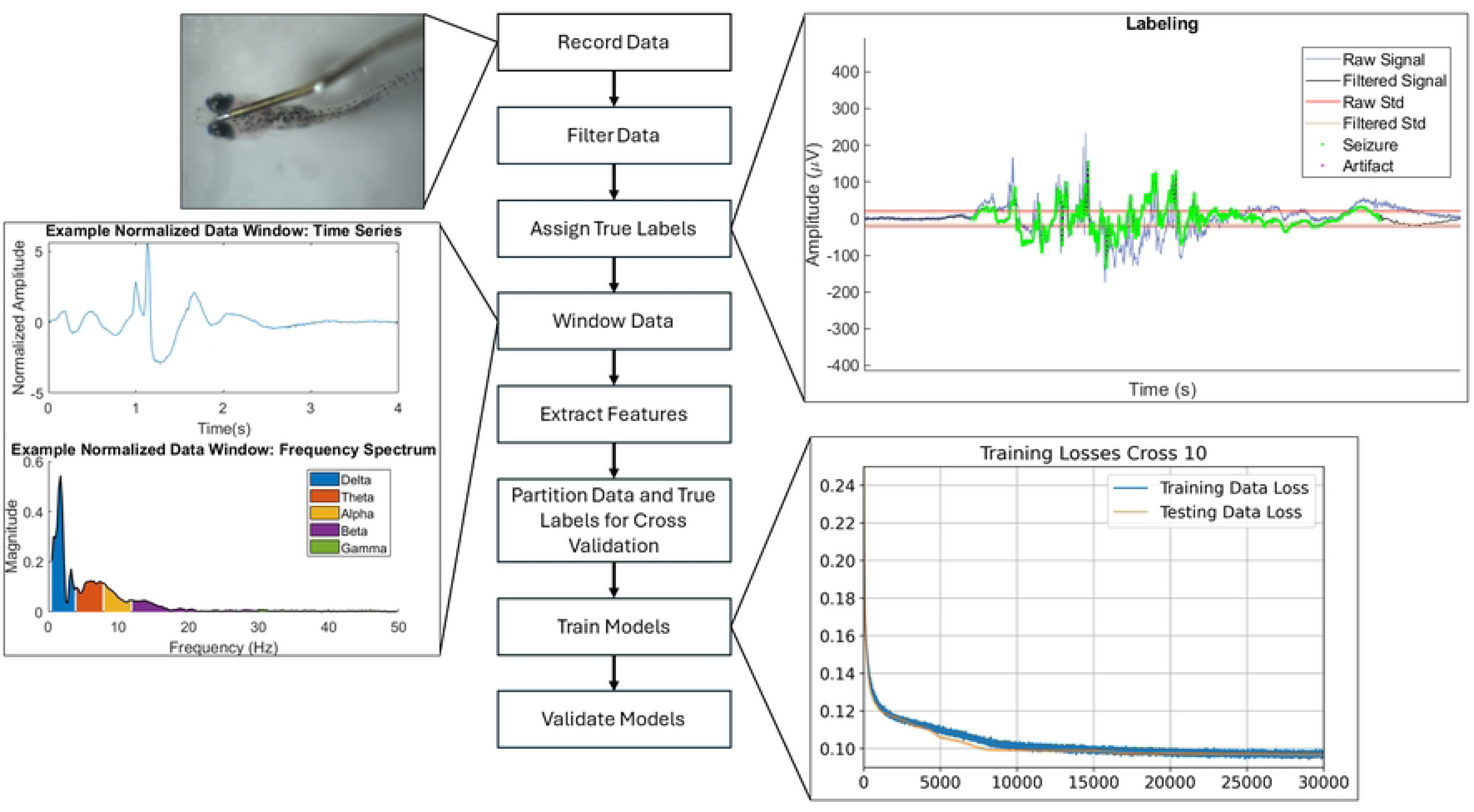
Methodology of Study. The workflow from raw recordings to collecting the algorithm training and validation results. Labels were generated using the filtered data overlaid on the raw recordings. Each cross-validation fold was class-balanced for both the training and validation data. The trained algorithms were utilized for validation and data excluded from the training process. The following results present the average accuracy of each algorithm across all cross-validation folds.

## Results

### Single Stage KNN Seizure Detection Algorithm

The average accuracy of the algorithm across all folds for all the K values tested can be seen in Fig 6 and Fig 7 for the 4-second and 1-second window data, respectively. The optimal K value was K=2 for the 4-second window data, with an 83.8% average accuracy on the validation data. Several example confusion matrices from this optimal K value are presented in Fig 8. The optimal K value for the 1-second window data was found to be K=2 as well and produced a validation accuracy of 82.8%. Example confusion matrices for these results are displayed in Fig 9. The trend among all cross-validation folds was for misclassification to occur primarily between the seizure and artifact classes.

**Fig 6.**
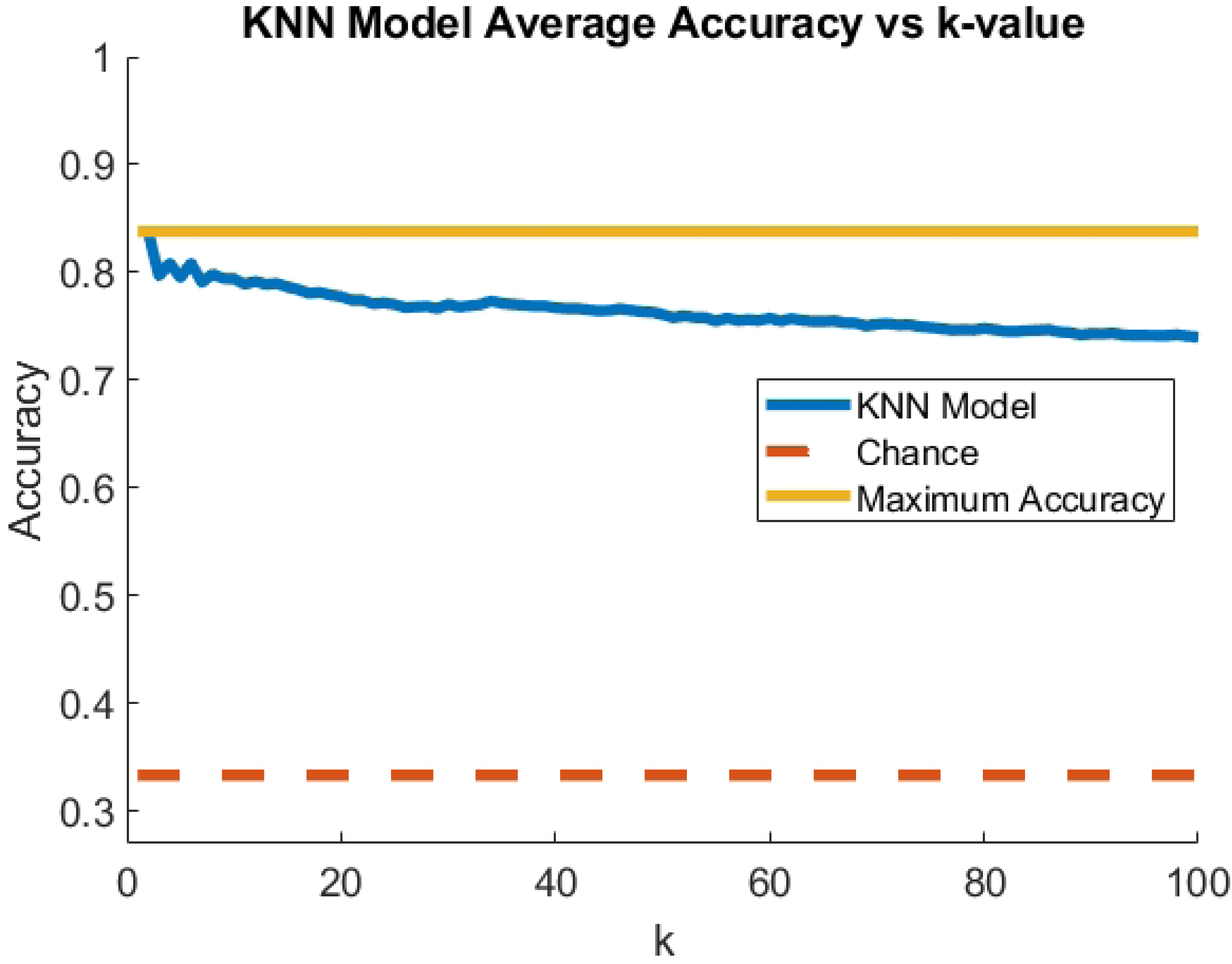
The plot of average accuracy across cross-validation fold for K values between 1 and 100 using 4-second windows. The optimal K value was determined to be K=2. The chance line is centered around 33% accuracy, as this detection task has three classes.

**Fig 7.**
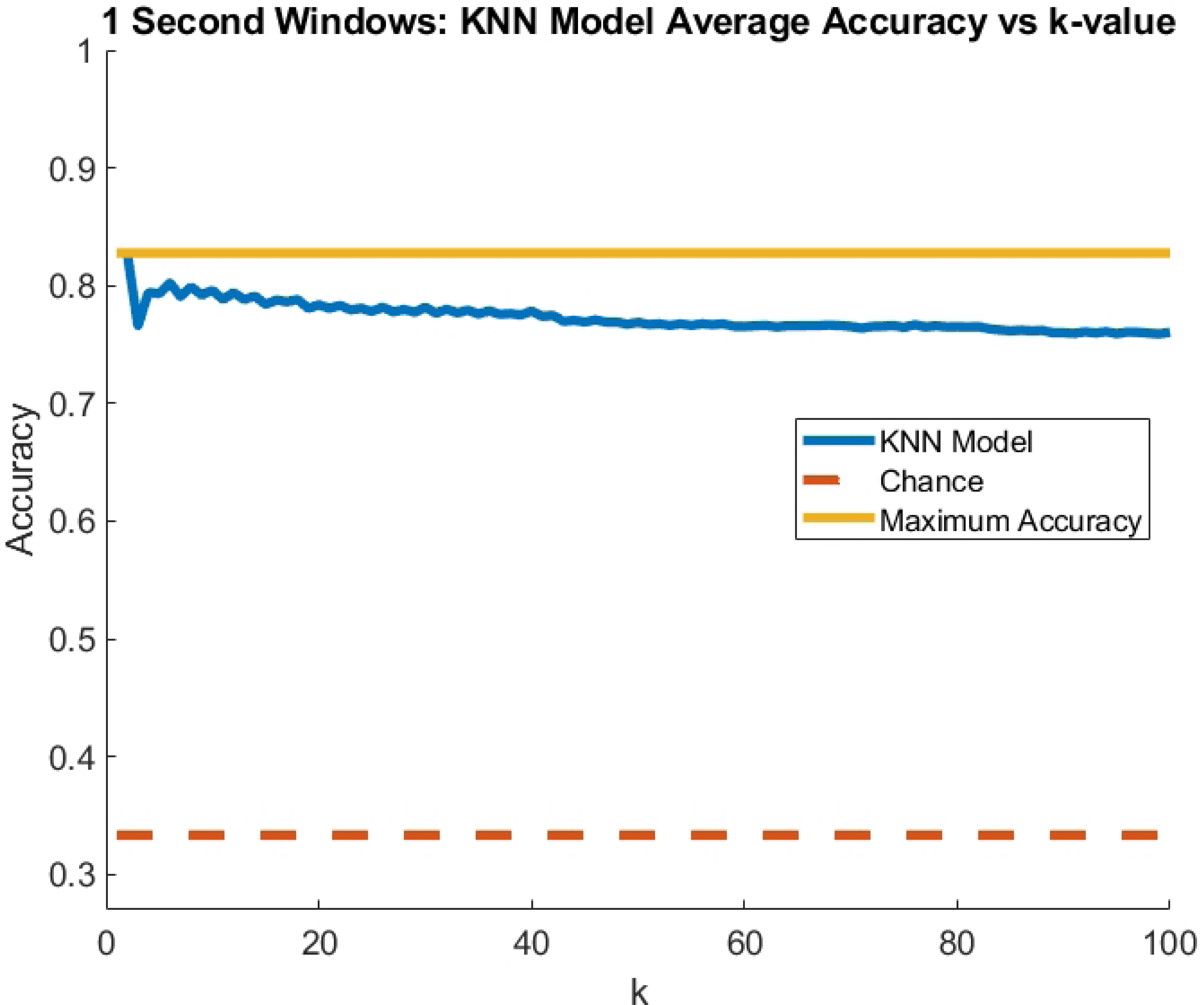
Plot of average accuracy across cross-validation fold for K values between 1 and 100 using 1-second windows. The optimal K value was determined to be K=2. The chance line is centered around 33% accuracy, as this detection task has three classes.

**Fig 8.**
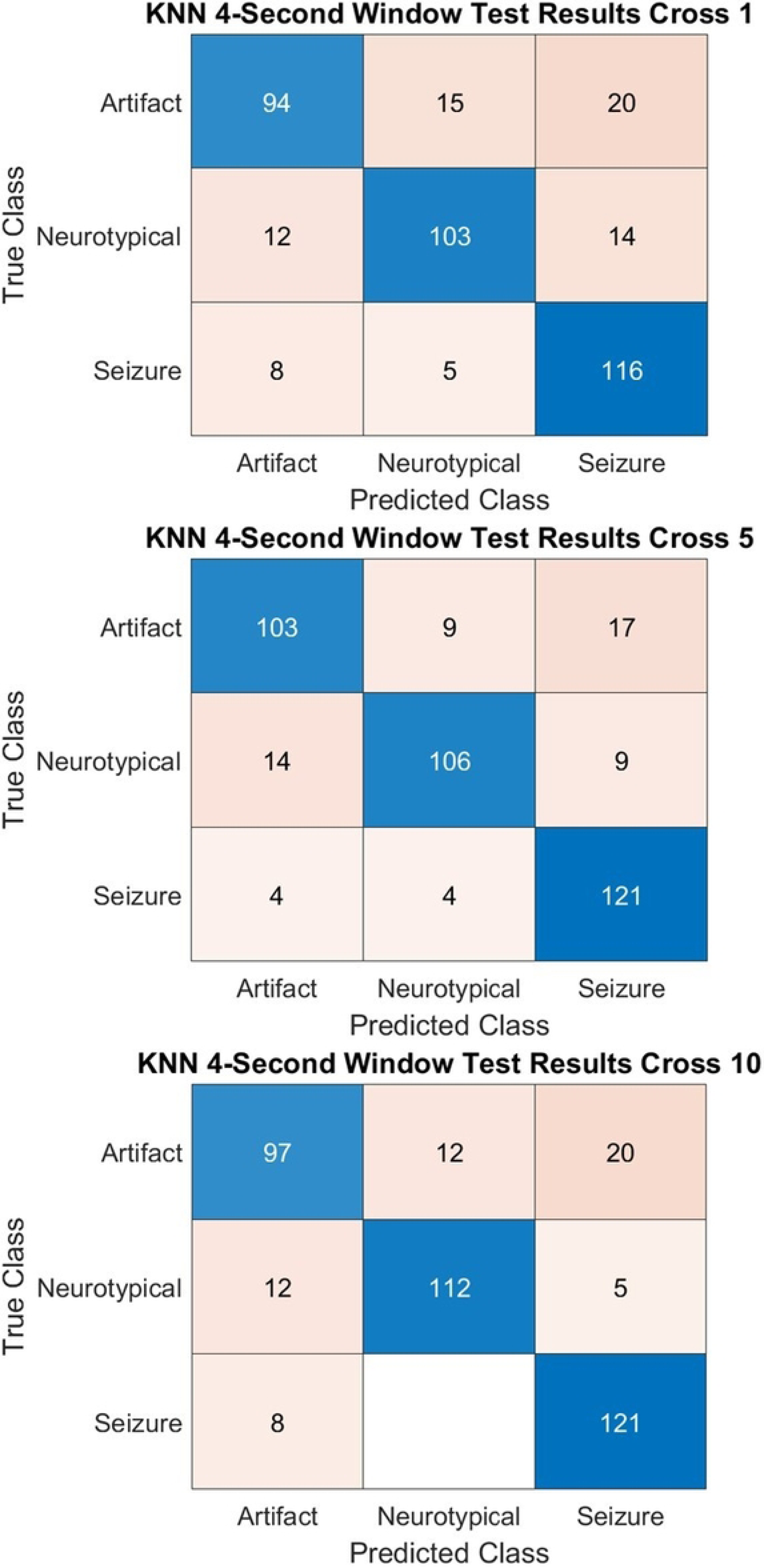
Example Single Stage KNN Confusion Matrices on 4-second window data. These confusion matrices characterize the results for three of the cross-validation folds from the Single Stage KNN detection algorithm on the validation data from that fold.

**Fig 9.**
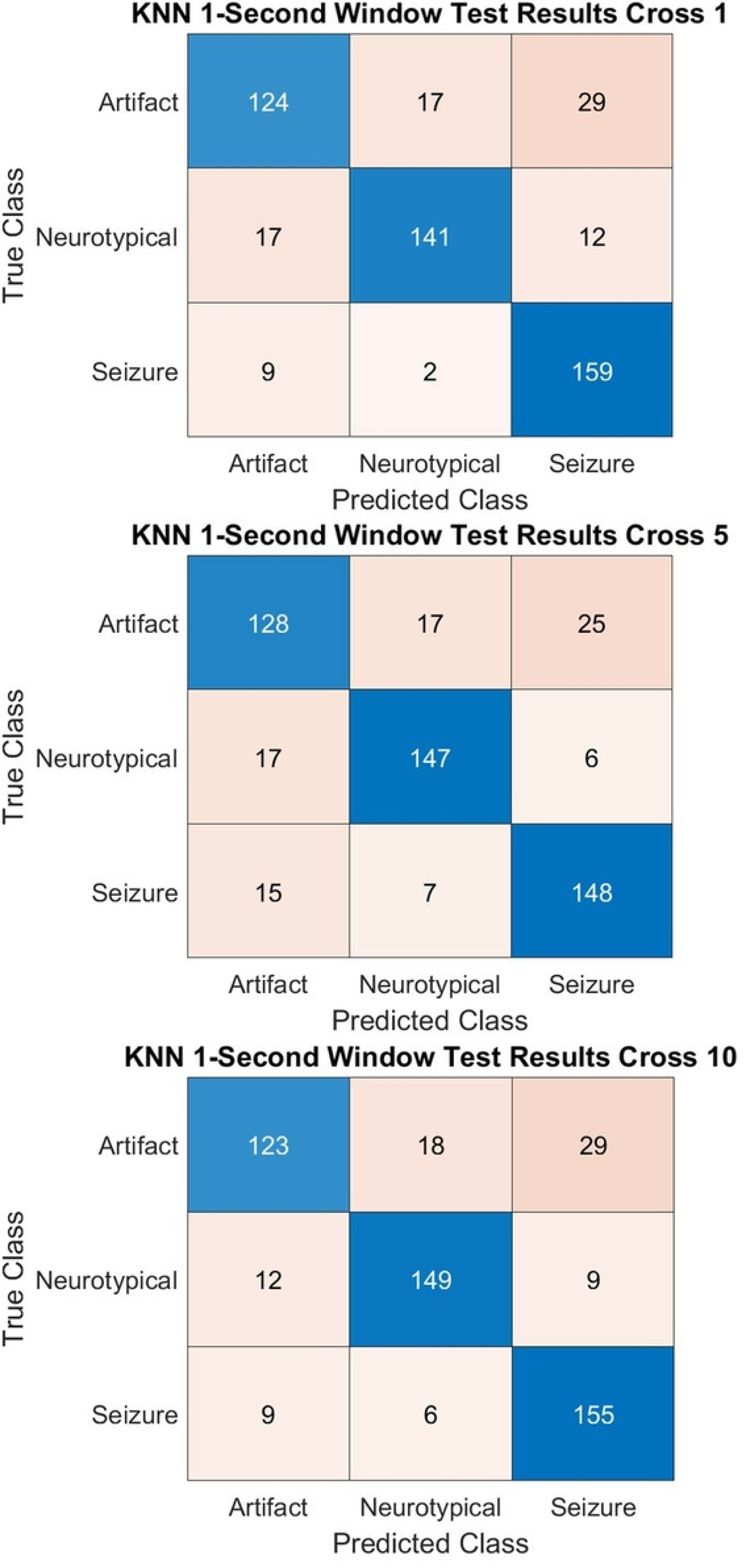
Example Single Stage KNN Confusion Matrices on 1-second window data. These confusion matrices characterize the results for three cross-validation folds from the Single Stage KNN detection algorithm on the validation data from that fold.

### Single Stage ANN Seizure Detection Algorithm

The average accuracy for the Single Stage ANN detection algorithm across all cross-validation folds for the 4-second window training data was 80.6%, and for the validation data from the same type of windows was 80.3%. The confusion matrices for the classification of the validation data for a subset of cross-validation folds are displayed in Fig 10. The average accuracy for the 1-second window data was 80.7% and 79.9% for the training and validation data, respectively. Example confusion matrices for these results are presented in Fig 11. The same trend as the Single Stage KNN algorithm occurred with the Single Stage ANN with more misclassifications between the seizure and artifact classes than any other classes.

**Fig 10.**
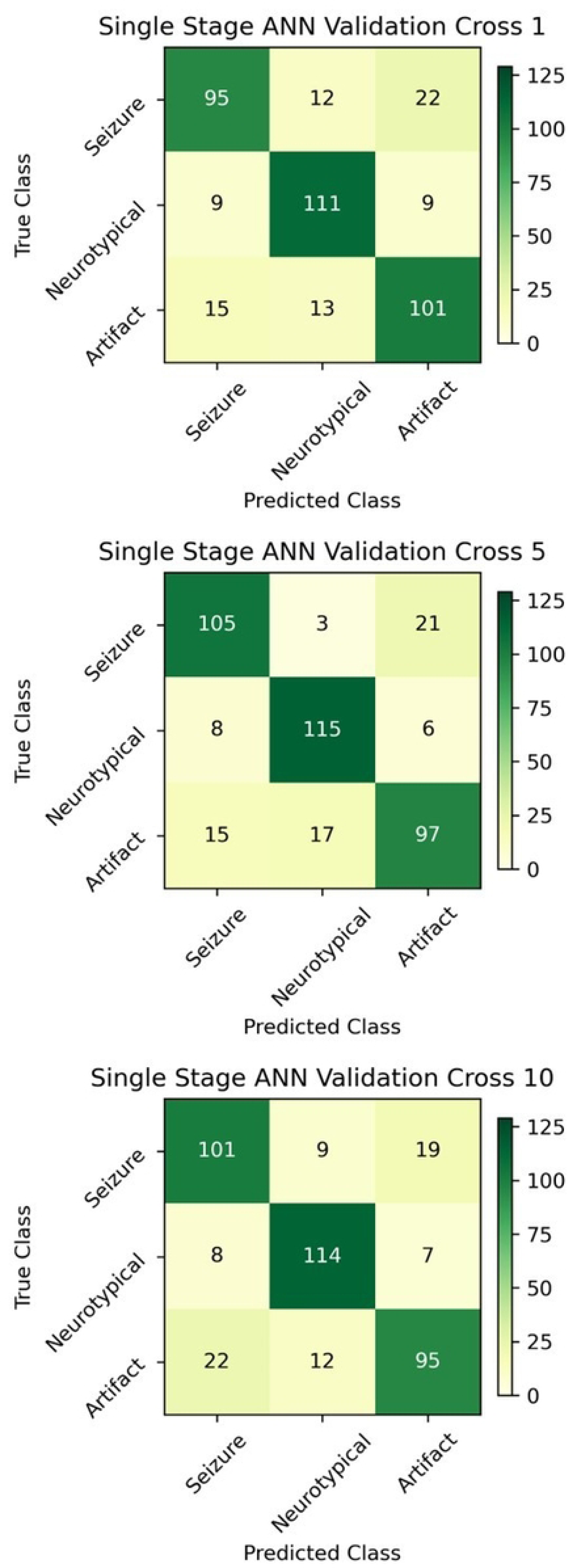
Example Single Stage ANN Confusion Matrices for 4 second windows. Confusion matrices for the validation data from a subset of cross-validation folds after training of the Single Stage ANN detection algorithm. The characteristics of these results hold for the other folds of the cross-validation, such as the tendency to misclassify between seizure and artifact.

**Fig 11.**
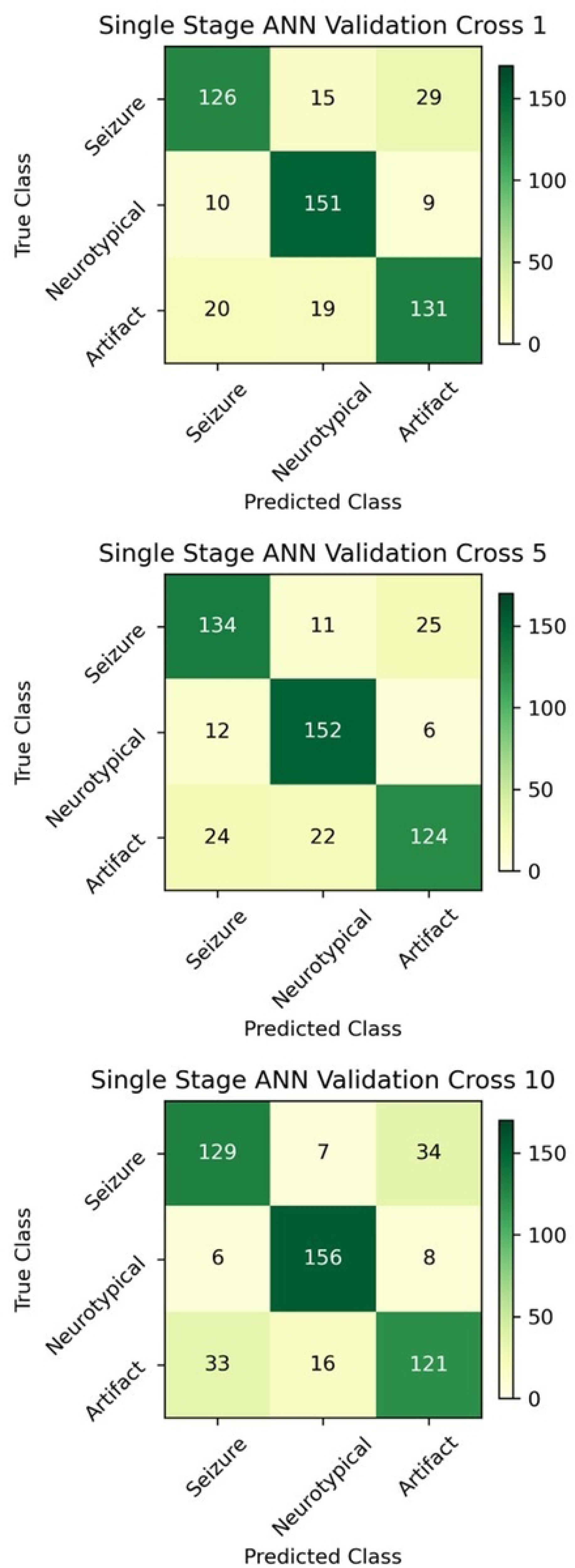
Example Single Stage ANN Confusion Matrices for 1-second windows. Confusion matrices for the validation data from a subset of cross-validation folds after training of the Single Stage ANN detection algorithm. The characteristics of these results hold for the other folds of the cross-validation, such as the tendency to misclassify between seizure and artifact.

### Two-Stage SVM Seizure Detection Algorithm

For optimizing the Two-stage SVM detection algorithm, the RBF kernel function was found to be the best performing out of all the kernels tested by a large margin for both 4- and 1- second windows. For the 4-second window data, the optimal gamma for the first stage was 0.1 and 0.2 for the second stage. The optimal C values for the first and second stages were 0.9 and 0.7, respectively. The average accuracy across cross-validation folds on the 4-second window data for the first stage was 98.0% and 89.3% for the training and validation data, respectively. The average accuracy for the second stage on the training data was 99.8%, and the validation data was 86.1%. When the stages were combined, accuracy for the training and validation data was 88.6% and 76.9%, respectively. Fig 12 displays confusion matrices for a representative subset of the cross-validation folds.

**Fig 12.**
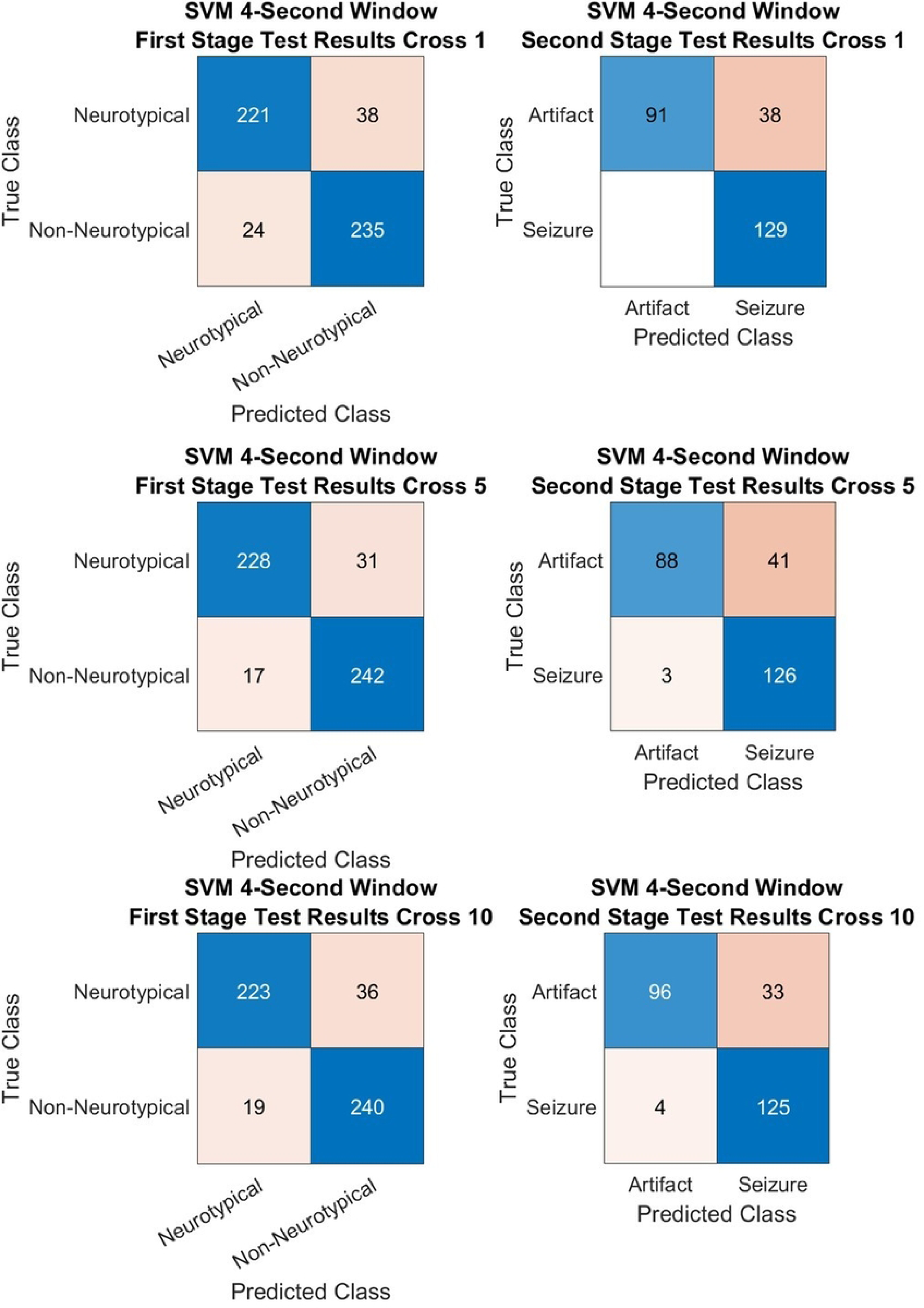
Example Two-Stage SVM Confusion Matrices for 4-second windows. These confusion matrices characterize the results for three of the cross-validation folds from the Two Stage SVM detection algorithm on the validation data from that fold. The second stage classification accuracy was worse than the first stage. This is expected as distinguishing neurotypical from non-neurotypical data is often easier than determining artifacts from seizure data.

When utilizing the 1-second window data, the optimal gamma values were found to be.95 for the first stage and .6 for the second stage. The optimal C value for the first stage was .1, and for the second stage was 1. The first stage average accuracy across cross-validation folds on the 1-second window data was 96.3% for the training data and 91.3% for the validation data. The average accuracy of the second stage for this same type of window was 98.5% and 86.6% for the training and validation, respectively. When both stages were combined, they achieved 90.4% average accuracy on the training data and 79.0% accuracy on the validation data. Example confusion matrices for this data are presented in Fig 13.

**Fig 13.**
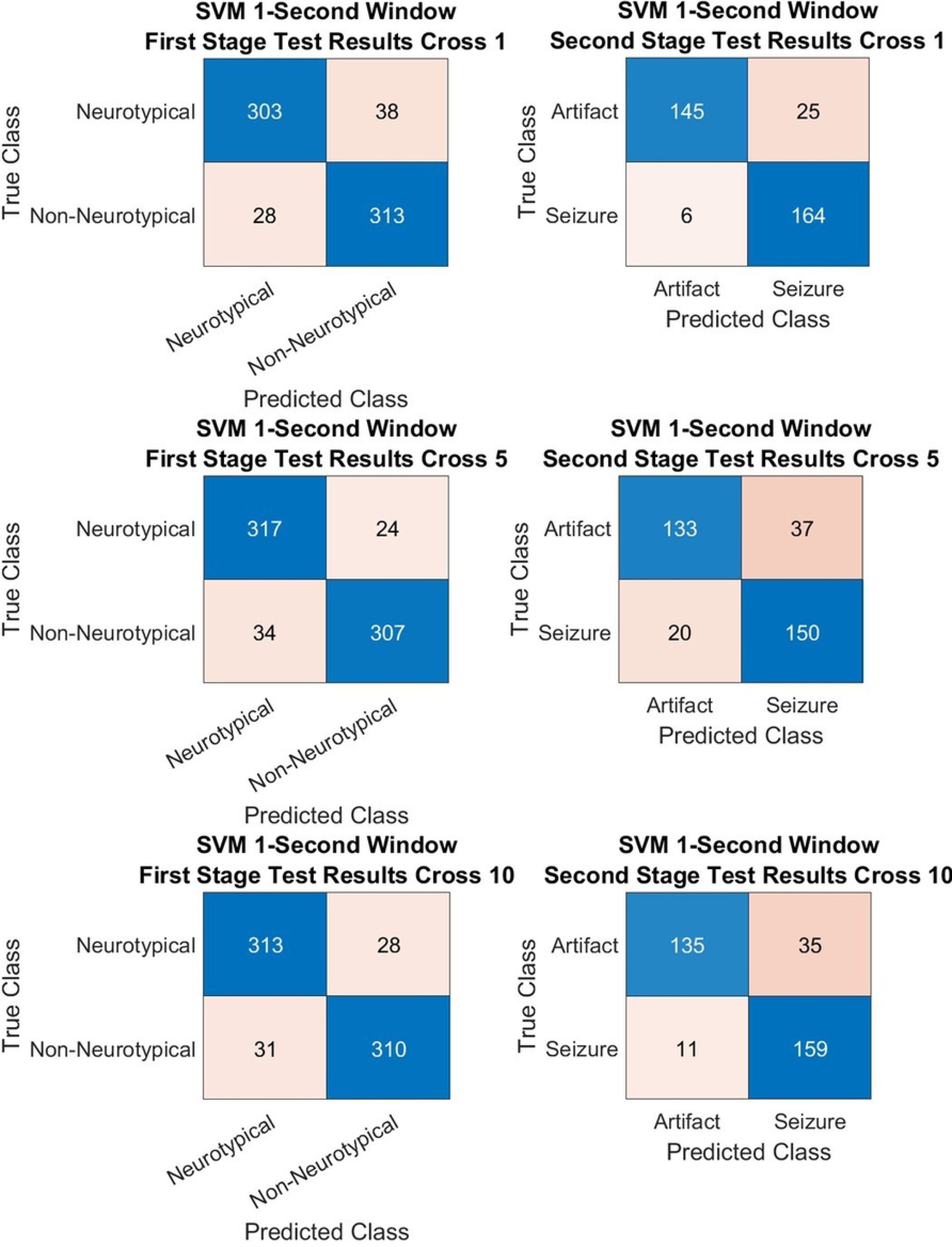
Example Two-Stage SVM Confusion Matrices for 1-second windows. These **c**onfusion matrices characterize the results for three of the cross-validation folds from the Two Stage SVM detection algorithm on the validation data from that fold. These confusion matrices demonstrate the same trend in performance between the first and second stages as the results from the 4-second windows.

### Two-Stage ANN Seizure Detection Algorithm

The Two Stage ANN detection algorithm achieved an average accuracy of 88.6% and 88.8% on the 4-second window training and validation data, respectively, with the first stage across all cross-validation folds. The average accuracy of the second stage on the same window type was 81.1% for the training data and 81.0% for the validation data. When being utilized in a combined fashion, they obtained a 71.9% average accuracy across cross-validation folds on both the training and validation data. A set of confusion matrices for a portion of the cross-validation folds is presented in Fig 14. This two-stage algorithm suffered from a similar issue as the Two-Stage SVM algorithm, as the second stage had reduced performance relative to the first stage.

**Fig 14.**
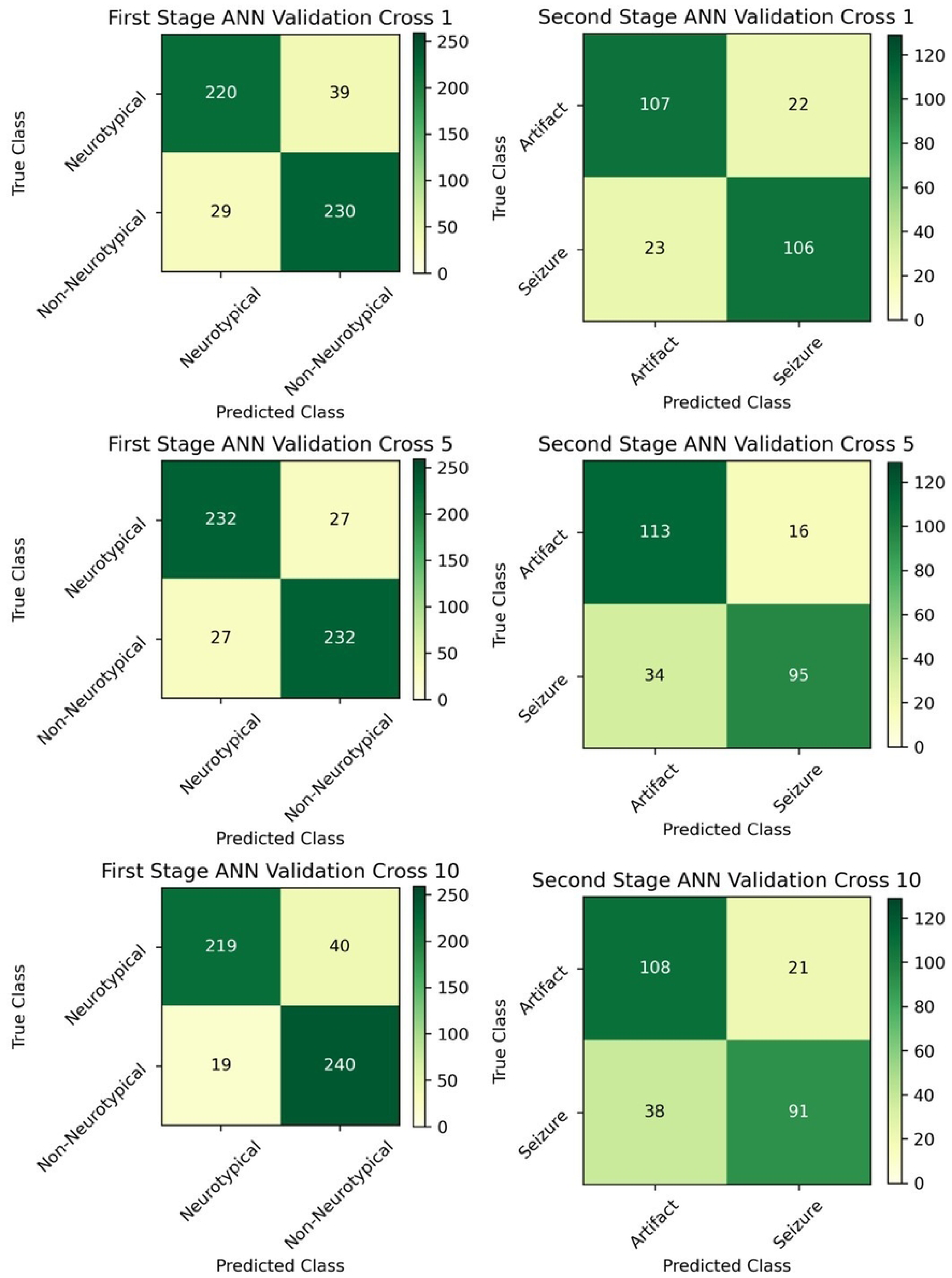
Example Two Stage ANN Confusion Matrices for 4-second windows. These confusion matrices characterize the results for three of the cross-validation folds from the Two Stage ANN detection algorithm on the validation data from that fold. The second stage classification accuracy is notably worse than the first stage. This pattern is the same as seen in the Two-stage SVM.

Utilizing the 1-second window, the first stage of Two-Stage ANN reached an accuracy of 90.4% and 91.0% on the training and validation data, respectively. The training accuracy of the second stage on the training data was 79.2%, and the validation data was 79.4%. When both stages were combined, they were only able to achieve an accuracy of 71.6% and 72.3% on the training and validation data, respectively. Example confusion matrices from individual folds of the cross-validation are presented in Fig 15.

**Fig 15.**
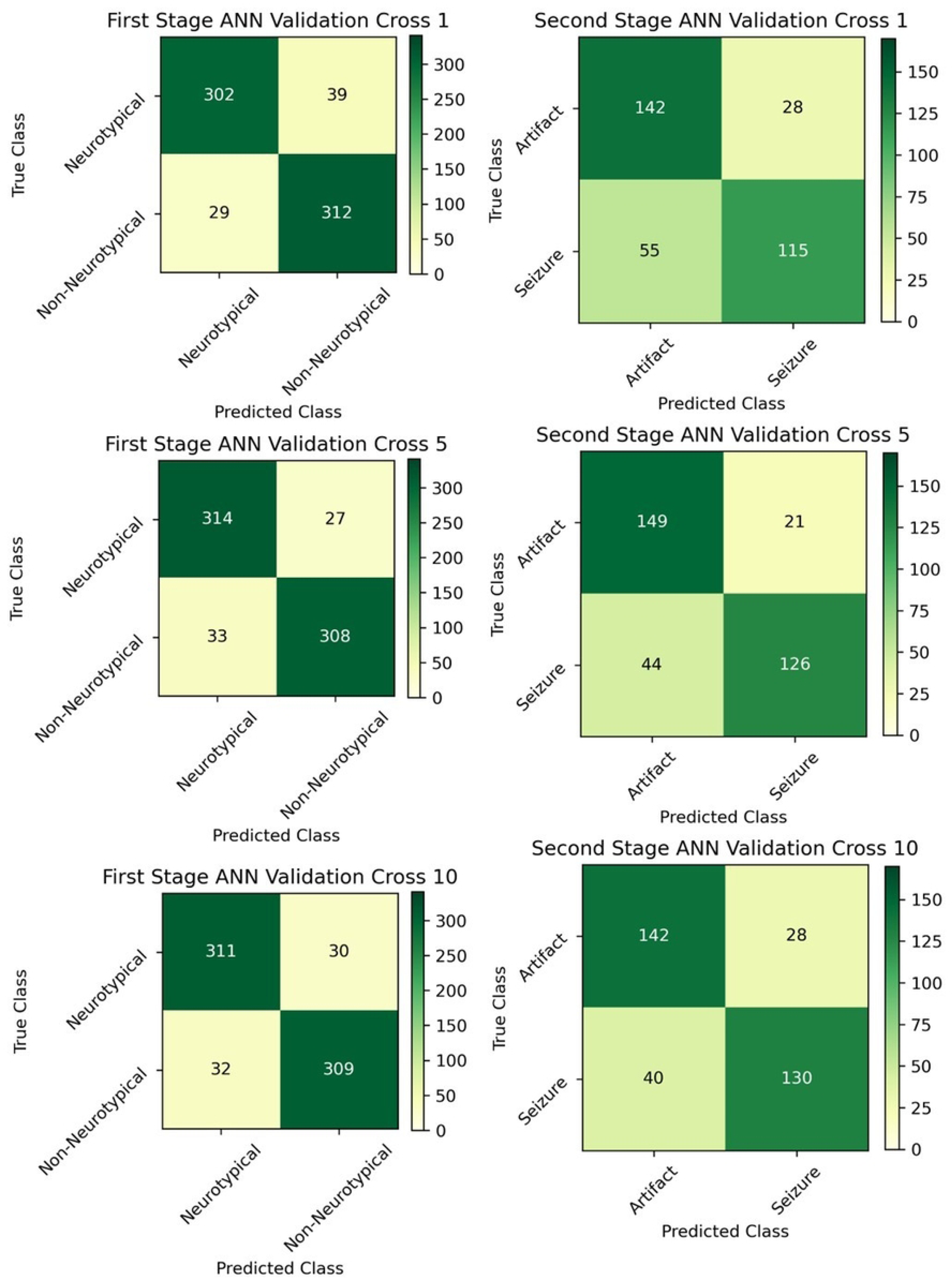
Example Two Stage ANN Confusion Matrices for 1-second windows. These confusion matrices characterize the results for three of the cross-validation folds from the Two Stage ANN detection algorithm on the validation data from that fold. The results follow the same trend as those from the 4-second windows.

A summary of all algorithm performances on training and validation data using 4-second windows with a 75% overlap is presented in Table 5. A summary of all algorithm performances on training and validation data using 1-second windows with a 50% overlap is presented in Table 6. Fig 16 displays the output from the four detection algorithms using 4-second windows from two segments of EEG data. Similarly, Fig 17 contains the detection algorithm output for the same two segments of EEG data but when using 1-second windows.

**Fig 16.**
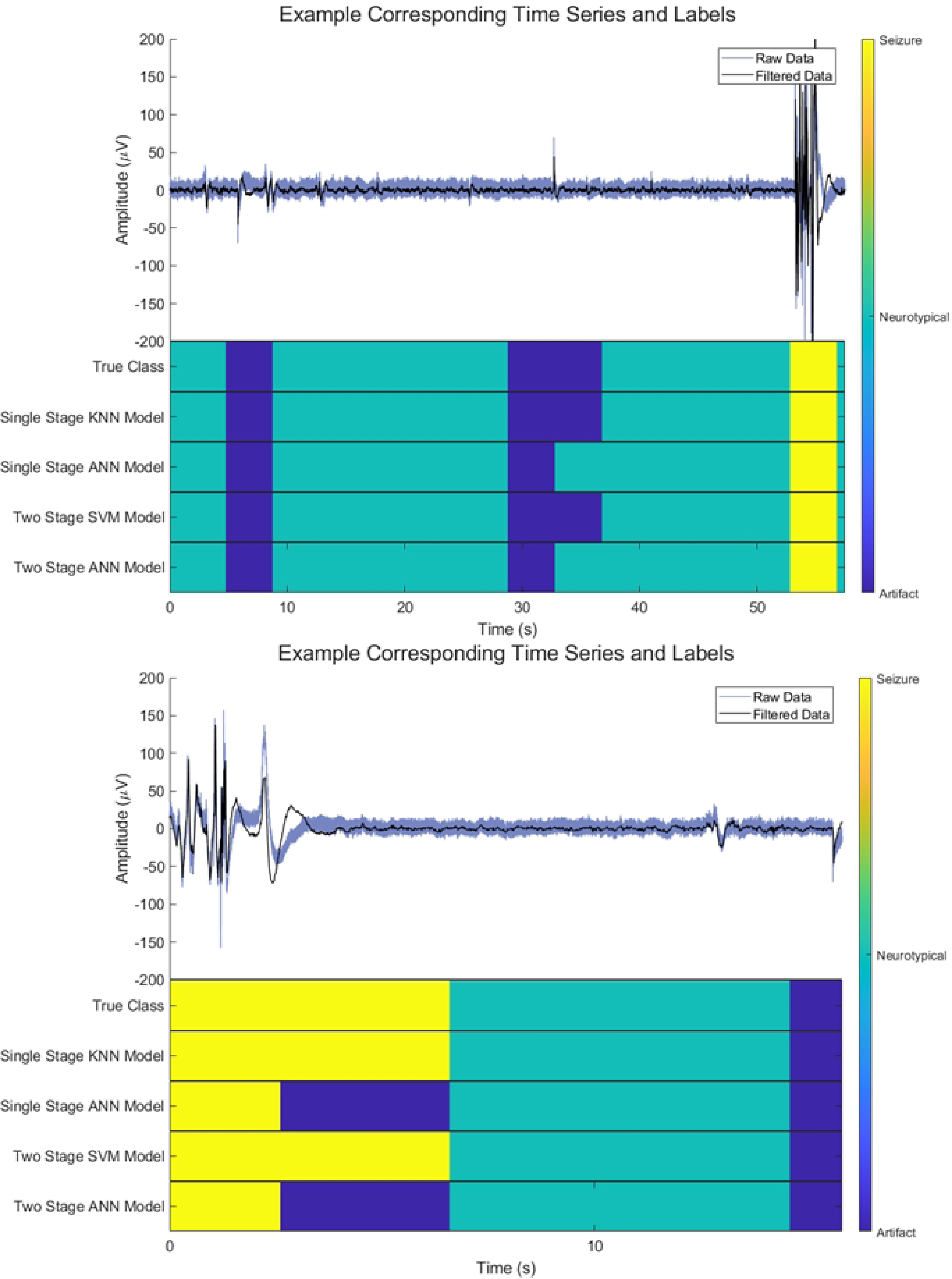
Example EEG Time Series and Corresponding Detection Algorithm Outputs using 4 Second Windows. Top) A segment of recorded EEG data passed through the detection algorithms and the respective output from each algorithm. Both the Single Stage and Two Stage ANN have classified the tail end of an impulse-like artifact as neurotypical. This is likely due to the amplitude of the artifact in this region being within a neurotypical range. Bottom) A shorter segment of recorded EEG data was passed through the detection algorithms and the respective output from each algorithm. Similar to the first example, the two ANN detection algorithms exhibit the same behavior; however, for this segment, they labeled the seizure’s tail end as an artifact. This is somewhat to be expected as the trend for the detection algorithms was to have the most misclassifications between seizures and artifacts.

**Fig 17.**
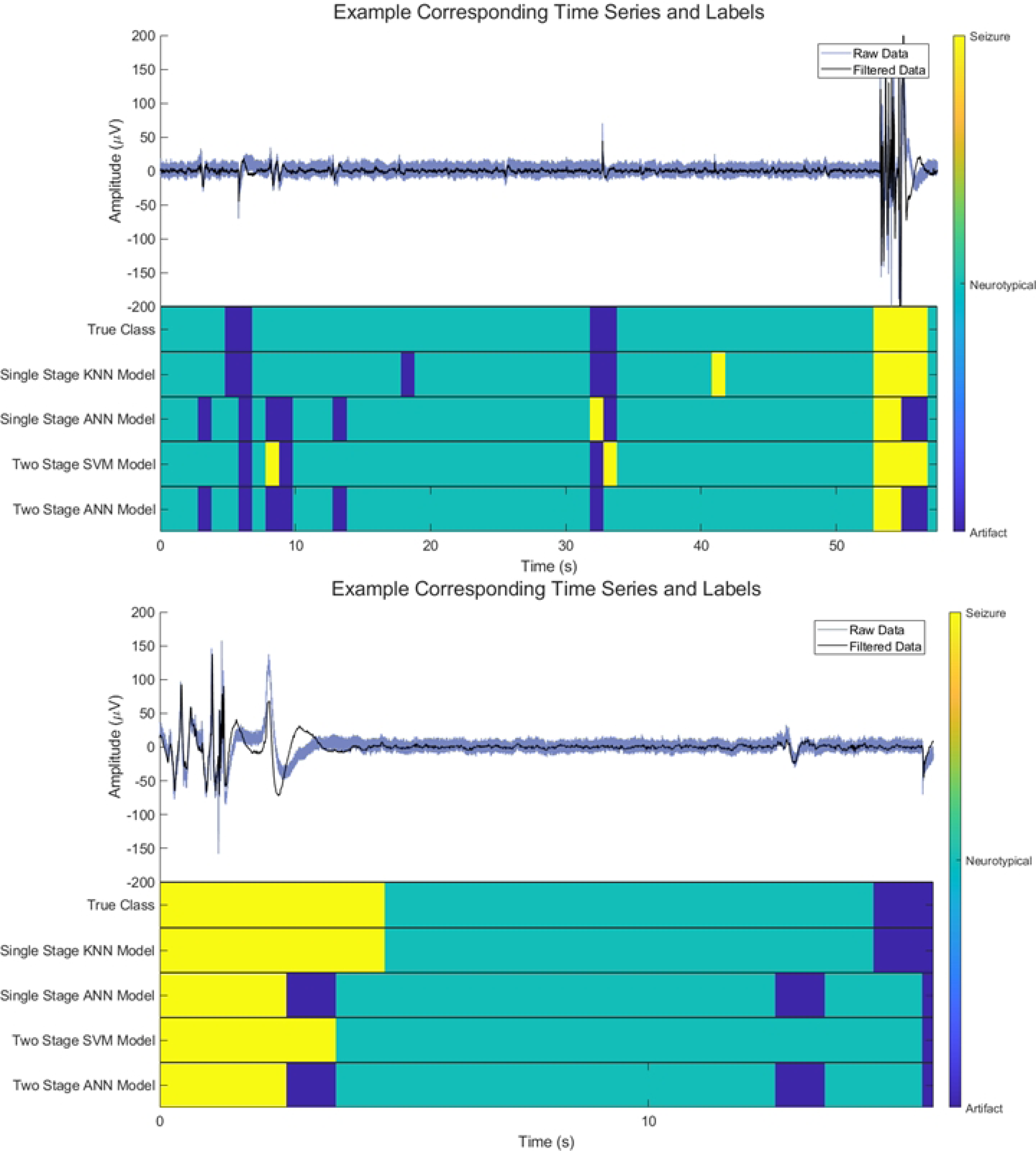
Example EEG Time Series and Corresponding Detection Algorithm Outputs using 1 Second Windows. Top) The same segment of recorded EEG data in the top plot of Fig 16 was passed through the detection algorithms and the respective output from each algorithm. Bottom) The same segment of recorded EEG data from the bottom plot of Figure 16 was passed through the detection algorithms and the respective output from each algorithm. The trend throughout all algorithms for both example segments was the classification of smaller changes in amplitude as non-neurotypical. This may suggest that the strict labeling scheme borrowed from Hunyadi *et al*. (2017) was not able to be accurately followed by the detection algorithms when using 1-second windows while at the same time decerning between seizure and artifact [18]. A less likely alternative is that their labeling procedure is too strict and misses some low amplitude artifacts.

**Table 5.**
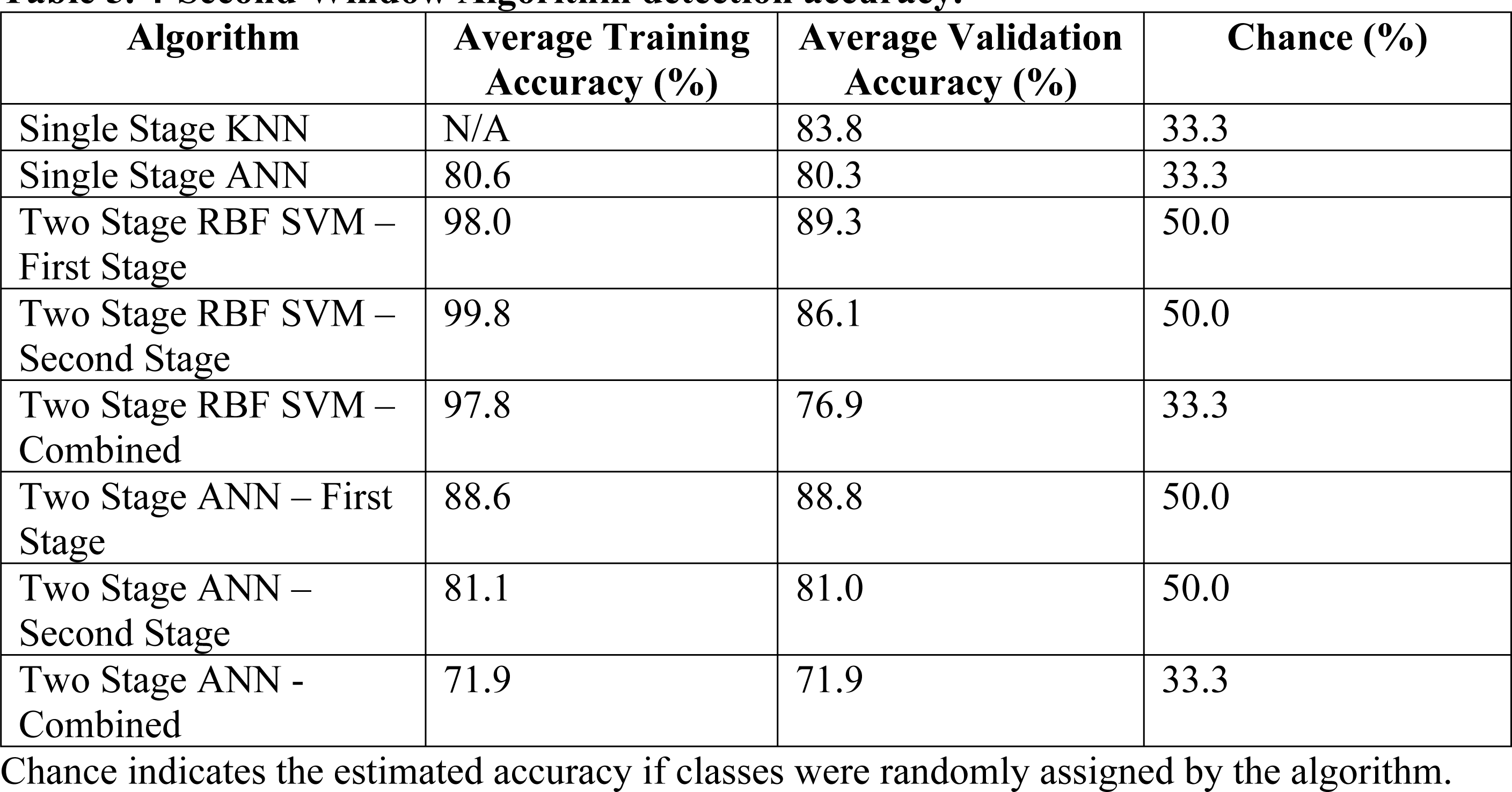
4-Second Window Algorithm detection accuracy.

**Table 6.**
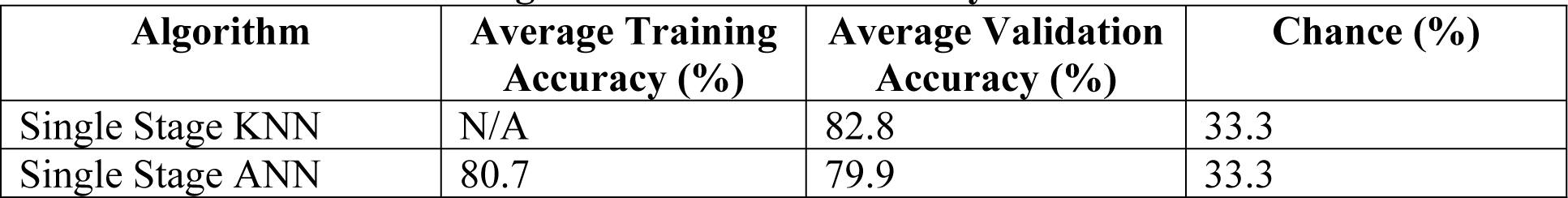
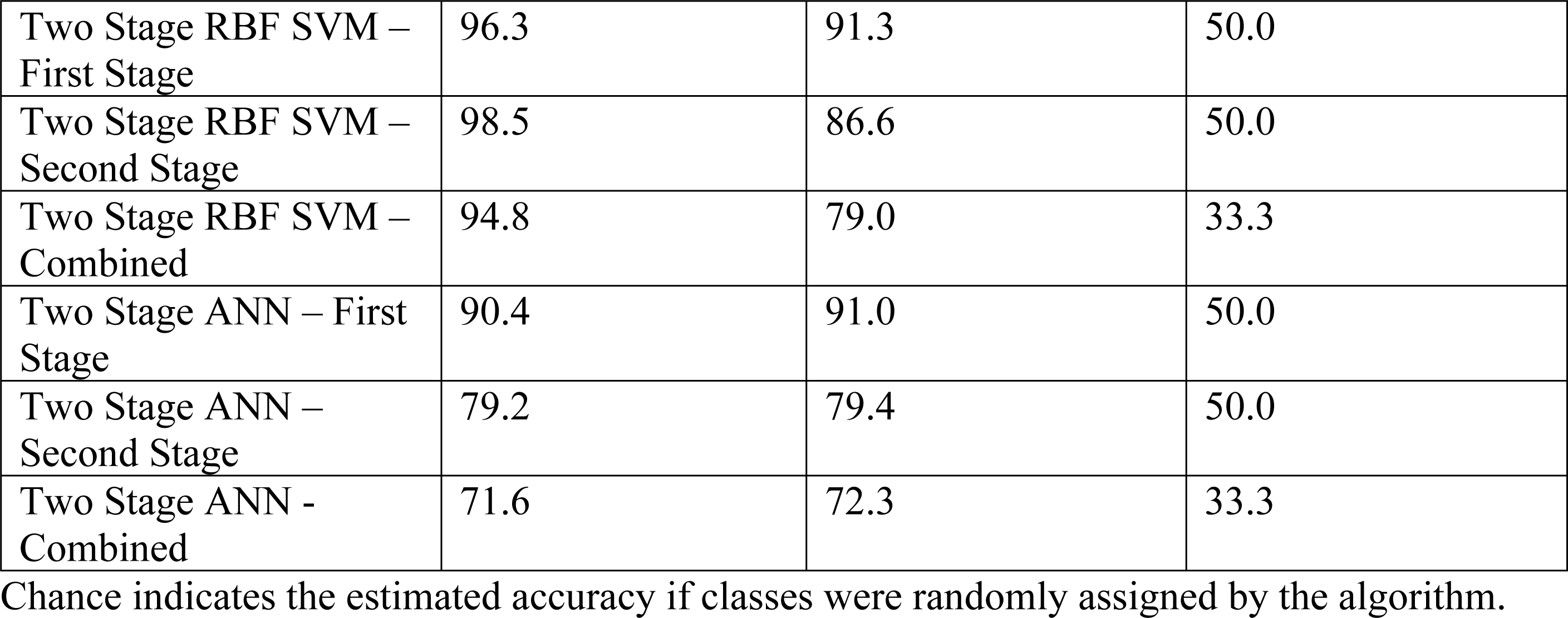
1-Second Window Algorithm detection accuracy.

| Algorithm | Average Training Accuracy (%) | Average Validation Accuracy (%) | Chance (%) |
| --- | --- | --- | --- |
| Single Stage KNN | N/A | 82.8 | 33.3 |
| Single Stage ANN | 80.7 | 79.9 | 33.3 |
| Two Stage RBF SVM – First Stage | 96.3 | 91.3 | 50.0 |
| Two Stage RBF SVM – Second Stage | 98.5 | 86.6 | 50.0 |
| Two Stage RBF SVM – Combined | 94.8 | 79.0 | 33.3 |
| Two Stage ANN – First Stage | 90.4 | 91.0 | 50.0 |
| Two Stage ANN – Second Stage | 79.2 | 79.4 | 50.0 |
| Two Stage ANN - Combined | 71.6 | 72.3 | 33.3 |

Chance indicates the estimated accuracy if classes were randomly assigned by the algorithm.

## Discussion

### Detection Algorithm Performance

The detection algorithm with the best performance overall on both types of windows was the Single Stage KNN. This is not unexpected as there is a precedent that KNN classifiers can achieve substantial results on seizure detection data [3]. Furthermore, the highly non-linear and inherent multiclass qualities of KNN predispose it to the present task of classifying data as seizure, artifact, or neurotypical.

The Single Stage ANN algorithm was the second-best performing algorithm for both types of windows. It was approximately 3% less accurate than the Single Stage KNN algorithm on the validation data across both types of windows. This may suggest that the single-stage algorithms offer some advantages over two-stage algorithms for this task. This could be due to bypassing errors compound from the first to the second stage by using multiclass classification. Additionally, the full power of an ANN detection algorithm may not be realized due to the limited size of the dataset.

The Two Stage SVM performed marginally better than the Two Stage ANN among the two-stage algorithms. This is primarily due to the second-stage ANN being slightly worse at accurately delineating seizures and artifacts from each other in comparison to the second-stage SVM. Both two-stage algorithms performed worse than the single-stage algorithms at the multiclass classification. However, the exceptionally high accuracy of the first stage of these algorithms may suggest that the two-stage algorithms provide some benefit over single-stage algorithms. Separating neurotypical data from non-neurotypical data could dramatically reduce the labeling time required by clinical technicians and researchers. One of these algorithms could be implemented to label neurotypical data automatically. Then, it could present only the non-neurotypical data to the technician, who could label it as a seizure or artifact. We believe that this technique could be implemented rapidly into existing labeling pipelines and dramatically reduce the time required to label individual files by technicians.

Comparing the results across both durations of windows, there appears to be a benefit to using the longer 4-second windows over the shorter 1-second windows for the single-stage algorithms. This may be because these algorithms may need more information to make an accurate classification than the 1-second window can afford. Interestingly, the two-stage algorithms performed better (except for the second stage of the Two-Stage ANN) using the 1- second window than using the 4-second window. These binary classifications are simpler components of the larger detection problem, which may have benefitted from having a larger dataset to train upon more than they were affected by the smaller duration of data producing to the extracted features.

Despite the Single Stage KNN detection algorithm being the best-performing, it could only achieve modest performance. This algorithm still suffered from the seizure-artifact misclassification issue that others displayed. The shared qualities between artifacts and seizures make seizure detection without prior artifact removal a complex challenge. Designing robust detection algorithms requires consideration of all possible features that may be able to delineate between artifact and seizure, incorporation of thorough preprocessing techniques to prepare the data for use properly, and careful algorithm implementation. Additionally, without a large, multi-individual dataset encompassing a wide variety of seizure types, obtaining higher classification accuracies may not be achievable.

### Limitations and Future Work

While this investigation serves as a strong starting point for the improvement of seizure detection algorithms, there are many facets of this work that limit the results. The foremost among these limitations was the small size of the dataset. We utilized just under 250 minutes of single-channel EEG data from zebrafish models in this investigation. This dataset used was primarily restricted due to a lack of labeled data. This issue is somewhat self-fulfilling as the overall goal of this project was to design algorithms to manage this task automatically. For the next stage of our investigation, more data will need to be labeled and added to the extant dataset.

Another limiting factor for our investigation was within the algorithms’ input structure.

The current iteration of these algorithms includes only 4 seconds or 1 second of data and none of the contextual information from the surrounding data before and after the recording. It is well established that EEG data can contain pre-ictal changes before a seizure occurs, which is why previous studies using human data have included larger windows of data than used in this investigation [3]. The obvious downside of these larger windows is the decreased resolution of the labeling procedure, which governed the decision to use a smaller window size for this investigation. A practical way to incorporate a larger span of time of data while also still having a small window size would be to use the features of data from several adjacent windows in addition to those in the primary window of concern. As this can rapidly increase the feature size of the dataset, a dimensionality reduction technique should be employed to reduce the feature set size if this strategy were to be used.

These two limitations of the current study are clear goals that should be targeted in future investigations. To reliably produce an accurate classification of epilepsy data, time must be spent increasing the quantity of accurately labeled data. Preprocessing, detection algorithms, and data structuring techniques must also be carefully implemented. The most challenging task facing automatic seizure detection algorithms is accurately labeling seizures and artifacts. This specific task should be a focus of future work, as improving this could lead to dramatically improved algorithm performance.

## Conclusion

We investigated improving automatic seizure detection algorithms for epilepsy research. This work specifically generated algorithms for use with zebrafish as a model organism for epilepsy. We developed four seizure detection algorithms: two single-stage architectures and two two-stage architectures. Among these algorithms, the Single Stage KNN utilizing 4-second duration windows performed best with a validation accuracy of 83.8% for multiclass classification of seizures, artifacts, and neurotypical data. Furthermore, the results from the two-stage classifiers indicate that implementing the first stage of these classifiers may benefit manual labeling by research and clinical technicians. The speed at which data could be labeled could be increased by only presenting technicians with the non-neurotypical data from a recording and automatically labeling neurotypical data. Results from this investigation serve as a launching point for further investigations into making improvements to seizure detection algorithms, which can improve the speed of labeling epilepsy EEG data for researchers and clinicians alike.

## Acknowledgments

We would like to acknowledge the assistance of Thales Guimaraes Parolari and Claudia Vianna Maurer-Morelli for their aid throughout the animal care and data collection process.

Furthermore, the authors would like to thank George Mason University and the University of Campinas for financial assistance and aid in housing the animals used in this study.

## Notes

### Competing Interest Statement

The authors have declared no competing interest.

## References

1. Engel J. ’’Surgery for seizures,’’ New Eng. J. Med. 1996; 334: 647–653.

2. Thurman DJ, Beghi E, Begley CE, Berg AT, Buchhalter JR, Ding D, Hesdorffer DC, Hauser WA, Kazis L, Kobau R., Kroner B, Labiner D, Liow K., Logroscino G, Medina MT. Standards for epidemiologic studies and surveillance of epilepsy. Epilepsia 2011; 7: 2–26.

3. Gotman J. Automatic recognition of epileptic seizures in the EEG. Electroencephalography and Clinical Neurophysiology. 1982; 54: Issue 5: 530–540. doi: 10.1016/0013-4694(82)90038-4.

4. Fergus P, Hignett D, Hussain A, Al-Jumeily D., Abdel-Aziz K. Automatic Epileptic Seizure Detection Using Scalp EEG and Advanced Artificial Intelligence Techniques. BioMed Research International. 2015. doi:10.1155/2015/986736

5. Kandratavicius L, Balista P, Lopes-Aguiar C, Ruggiero R, Umeoka E, Garcia-Cairasco N, Bueno-Junior L, Leite J. Animal models of epilepsy: use and limitations. Neuropsychiatr Dis Treat. 2014;10:1693–1705. doi: 10.2147/NDT.S50371

6. Afrikanova T, Serruys ASK, Buenafe OEM, Clinckers R, Smolders I, de Witte PAM, Crawford AD, Esguerra CV. Validation of the zebrafish pentylenetetrazol seizure model: loco motor versus electrographic responses to antiepileptic drugs. PLoS One 2013; 8:1–8. doi: 0.1371/journal.pone.0054166 2.

7. Howe K, Clark MD, Torroja CF, Torrance J, Berthelot C, Muffato M, et al. The zebrafish reference genome sequence and its relationship to the human genome. Nature. 2013; 496:498–503. doi: 10.1038/nature12111.

8. Lieschke GJ, Currie PD. Animal models of human disease: zebrafish swim into view. Nat Rev Genet. 2007; 8: 353–367. doi: 10.1038/nrg2091

9. Zheng J, Hsieh F, Ge L. A Data-Driven Approach to Predict and Classify Epileptic Seizures from Brain-Wide Calcium Imaging Video Data. IEEE/ACM Transactions on Computational Biology and Bioinformatics. 2020. 17: 1858–1870. doi: 10.1109/TCBB.2019.2895077.

10. Burrows DRW, Samarut É, Liu J., Baraban SC, Richardson MP, Meyer MP, Rosch RE. Imaging epilepsy in larval zebrafish. European Journal of Paediatric Neurology. 2020; 24: 70–80. doi: 10.1016/j.ejpn.2020.01.006.

11. Cho SJ, Byun D, Nam TS, Choi SY, Lee BG, Kim MK, Kim S. Zebrafish as an animal model in epilepsy studies with multichannel EEG recordings. Sci Rep. 2017; 7: 3099. doi: 10.1038/s41598-017-03482-6

12. Zdebik AA, Mahmood F, Stanescu HC, Kleta R, Bockenhauer D, Russell C. Epilepsy in kcnj10 Morphant Zebrafish Assessed with a Novel Method for Long-Term EEG Recordings. PLoS ONE. 2013; 8(11). doi: 10.1371/journal.pone.0079765

13. Meyer M, Dhamne SC, LaCoursiere CM, Tambunan D, Poduri A, Rotenberg A. Microarray Noninvasive Neuronal Seizure Recordings from Intact Larval Zebrafish. PLoS ONE. 2016; 11(6). doi: 10.1371/journal.pone.0156498

14. Hong S, Lee P, Baraban SC, Lee L. A Novel Long-term, Multi-Channel and Non-invasive Electrophysiology Platform for Zebrafish. Sci Rep. 2016; 6: 28248. doi: 10.1038/srep28248

15. Cho SJ, Nam TS, Choi SY, Kim MK, and Kim S, 3D printed multi-channel EEG sensors for zebrafish. IEEE SENSORS. 2015; 1–3. doi: 10.1109/ICSENS.2015.7370544.

16. Lee Y, Lee KJ, Jang JW, Lee S, Kim S. An EEG system to detect brain signals from multiple adult zebrafish. Biosensors and Bioelectronics. 2020; 164. doi: 10.1016/j.bios.2020.112315.

17. Milder PC, Zybura AS, Cummins TR, Marrs JA. Neural Activity Correlates With Behavior Effects of Anti-Seizure Drugs Efficacy Using the Zebrafish Pentylenetetrazol Seizure Model. Frontiers in Pharmacology. 2022; 13. doi: 10.3389/fphar.2022.836573

18. Hunyadi B, Siekierska A, Sourbron J, Copmans D, de Witte PAM. Automated analysis of brain activity for seizure detection in zebrafish models of epilepsy. Journal of Neuroscience Methods. 2017; 287: 13–24. doi: 10.1016/j.jneumeth.2017.05.024.

19. Xing Q, Huynh V, Parolari TG, Maurer-Morelli CV, Peixoto N, Wei Q. Zebrafish larvae heartbeat detection from body deformation in low resolution and low frequency video. Med Biol Eng Comput. 2018; 56: 2353–2365. doi: 10.1007/s11517-018-1863-7

20. Baraban SC, Taylor MR, Castro PA, Baier H. Pentylenetetrazole induced changes in zebrafish behavior, neural activity and c-fos expression. Neuroscience. 2005; 131(3): 759–768. doi: 10.1016/j.neuroscience.2004.11.031.

21. Mussulini BHM, Leite CE, Zenki KC, Moro L, Baggio S, Rico EP, et al. Seizures Induced by Pentylenetetrazole in the Adult Zebrafish: A Detailed Behavioral Characterization. PLoS ONE. 2013; 8(1). doi: 10.1371/journal.pone.0054515

22. Pineda R, Beattie CE, Hall CW. Recording the adult zebrafish cerebral field potential during pentylenetetrazole seizures. Journal of Neuroscience Methods. 2011; 200(1): 20–28. doi: 10.1016/j.jneumeth.2011.06.001.

